# 2000-year fish bone record reveals transition to commercial fisheries during climatic change

**DOI:** 10.1101/2025.10.14.682357

**Authors:** Danielle L. Buss, Abigail K. Parker, Mohsen Falahati-Anbaran, Indrė Žliobaitė, Rory Connolly, Thomas C.A. Royle, Rachel Ballantyne, Adam Boethius, Monica K. Dütting, Monica Nordanger Enehaug, Inge Bødker Enghoff, Anton Ervynck, Sheila Hamilton-Dyer, Jennifer F. Harland, Richard C. Hoffmann, Poul Holm, Anne Karin Hufthammer, Inge van der Jagt, Beatrice Krooks, Hans Christian Küchelmann, Fredrik Charpentierppl Ljungqvist, Lembi Lõugas, Ola Magnell, Daniel Makowiecki, Emma Maltin, Hanneke J.M. Meijer, William F. Mills, Rebecca Nicholson, Liz Quinlan, Hannah Russ, Kenneth Ritchie, Andrea Seim, Wim Van Neer, Wim Wouters, James H. Barrett

**Affiliations:** Department of Archaeology and Cultural History, NTNU Vitenskapsmuseet, Norwegian University of Science and Technology, Trondheim, Norway; Department of Computer Science; Department of Geosciences and Geography, University of Helsinki, Helsinki, Finland; Trinity Centre for Environmental Humanities, Trinity College Dublin, Dublin, Ireland; McDonald Institute for Archaeological Research, University of Cambridge, Downing Street, Cambridge, CB2 3ER, UK; Department of Archaeology and Ancient History, University of Lund, Lund, Sweden; Zaandam, The Netherlands; Kirkehøj 6, DK-2900, Hellerup, Denmark; Flemish Heritage Agency, Brussels, Belgium; SH-D ArchaeoZoology, Southampton, UK; Archaeology Institute, University of the Highlands and Islands, Orkney College, Kirkwall, Orkney, UK; Department of Archaeology, University of York, Heslington, York, UK; Department of History, York University, Toronto, Ontario, Canada; Department of Natural History, University Museum of Bergen, University of Bergen, Bergen, Norway; Department of Archaeology, Cultural Heritage Agency of the Netherlands, Amersfoort, Netherlands; Department of Archaeology, Ancient History and Conservation, Uppsala University - Campus Gotland, Sweden; German Maritime Museum, Leibniz Institute for German Maritime History, Hans-Scharoun-Platz-1, Germany; Department of History, Stockholm University, Stockholm, Sweden; Bolin Centre for Climate Research, Stockholm University, Stockholm, Sweden; Archaeological Research Collection, Tallinn University, Tallinn, Estonia; The Archaeologists, National Historical Museums, Lund, Sweden; Department of Environmental Archaeology and Human Palaeoecology, Institute of Archaeology, Faculty of History, Nicolaus Copernicus University, Toruń, Poland; Department of Archaeology and Classical Studies, Stockholm University, Stockholm, Sweden; Bohusläns museum, Uddevalla, Sweden; Department of Geography and Environmental Science, University of Reading, Reading, UK; Oxford Archaeology, Osney Mead, Oxford, UK; Archaeology.biz, Barnard Castle, UK; Moesgaard Museum, Aarhus University, Moesgaard, Denmark; Institute of Forest Sciences, University of Freiburg, Freiburg, Germany; Department of Botany, University of Innsbruck, Innsbruck, Austria; Regional Climate Group, Department of Earth Science, University of Gothenburg, Gothenburg, Sweden; Royal Belgian Institute of Natural Sciences, Brussels, Belgium; Laboratory of Biodiversity and Evolutionary Genomics, University of Leuven; Leuven, Belgium

## Abstract

Animal bones from archaeological contexts can reveal the interplay between past environments and human societies. Resource acquisition shaped many aspects of past societies and influenced the development of trade networks and migration. Fish have been a cornerstone of human subsistence for millennia, yet the rise of commercial fishing and trade was complex. Here, we synthesised a database of ∼1.9 million zooarchaeological fish records spanning 2000 years across Europe. Using machine-learning of catch compositions alongside fish thermal tolerances, we show that fisheries became less local over time, with homogenisation coinciding with Little Ice Age-associated cooling, a period of documented resource scarcity, concurring with growing trade. Moreover, increased proportions of marine taxa and more specialist marine fisheries were observed in the preceding Medieval Climate Anomaly, to sustain concurrent urban and population growth. Enhanced use of marine protein buffered food insecurity, whilst signalling the transition from localised to trans-regional trade networks.

## Introduction

Fish have been harvested for tens of thousands of years^1^, with early innovations in fishing and preservation making them one of the first globally traded food commodities^2,3^, and remain a key protein source for billions of people today^4^. As a result of this long-standing reliance, changes in fish stock availability have shaped human demographics and cultural practices worldwide for millennia, with a dynamic interplay between thriving coastal, lacustrine, and riparian communities resulting in demographic growth, and diminished or locally insufficient resources resulting in depopulation, or the adoption of novel fishing strategies^5–8^.

Periods of resource scarcity have historically driven the expansion of trade networks, enabling communities to source goods from unaffected regions and gain knowledge to facilitate adaptation to environmental stressors^9,10^. The emergence of long-range trade networks has been closely linked to migration events and urbanisation^11,12^, especially for fisheries^8,13,14^. While the emergence of trans-regional European fisheries has been discussed qualitatively within these multicausal frameworks^8,15–17^, a comprehensive, data-driven, across-border analysis of these transitions, particularly for periods lacking systematic historical records, remains absent. To address this gap and elucidate the chronology and potential causation of the emergence of trans-regional fisheries, we apply a novel machine-learning approach to synthesise archaeological fish bone from archaeological contexts that indicate harvesting, storage and/or consumption by humans.

Past responses to changing resource availability can be traced through zooarchaeological time-series, including excavated bones, shells, and otoliths, which provide indications of past harvesting practices as well as environmental and ecological change^18^. For instance, zooarchaeological and palaeontological evidence has revealed animal extinctions (e.g. gray whales (*Eschrichtius robustus*) in the North Atlantic and the Waitaha penguin (*Megadyptes waitaha*) in New Zealand^19,20^) as well as declines in species abundance within particular regions (e.g. Atlantic bluefin tuna (*Thunnus thynnus*) in the Mediterranean, Atlantic salmon (*Salmo salar*) in the Palaeo-Rhine catchment, and North Atlantic right whales (*Eubalaena glacialis*) in Europe^21–23)^. Conversely, species range expansions have been driven by aquaculture use (e.g. carp, *Cyprinus carpio*^17^). Moreover, changes in the community structure of species characteristic of specific temperature gradients can indicate past climatic shifts and concomitant shifts in the fish resource base^24,25^.

By analysing life history traits and thermal tolerances from a database of more than 1.9 million archaeological fish bone records from Europe dating between 1 CE and 2000 CE (**Fig. 1**), we demonstrate a homogenisation of fish consumption between 1100 – 1700 CE, together with a trans-regional standardisation in reliance on marine resources during the Medieval Climate Anomaly (MCA; also referred to as the Medieval Warm Period, c. 950 to 1250 CE^26^; a time of high terrestrial air temperatures and human population increase, alongside decreasing sea surface temperatures in oceanic and coastal regions). We further identify a propensity towards marine-only fisheries starting in the MCA and continuing across much of the Little Ice Age (c. 1300 to 1850 CE), a period characterised by colder terrestrial climatic conditions and reduced agricultural production in parts of Europe^26–29^. The results indicate that the reliance on marine taxa was associated with a shift from localised to more consistent trans-regional fishing practices within Europe and with long-range exchange networks established during the MCA and Little Ice Age. Fish bone records are provided through the open-access Northern Seas Synthesis Database (NSS-DB), which represents novel insights into fishing pressures and human consumption of aquatic resources over the past two millennia across Europe.

**Figure 1.**
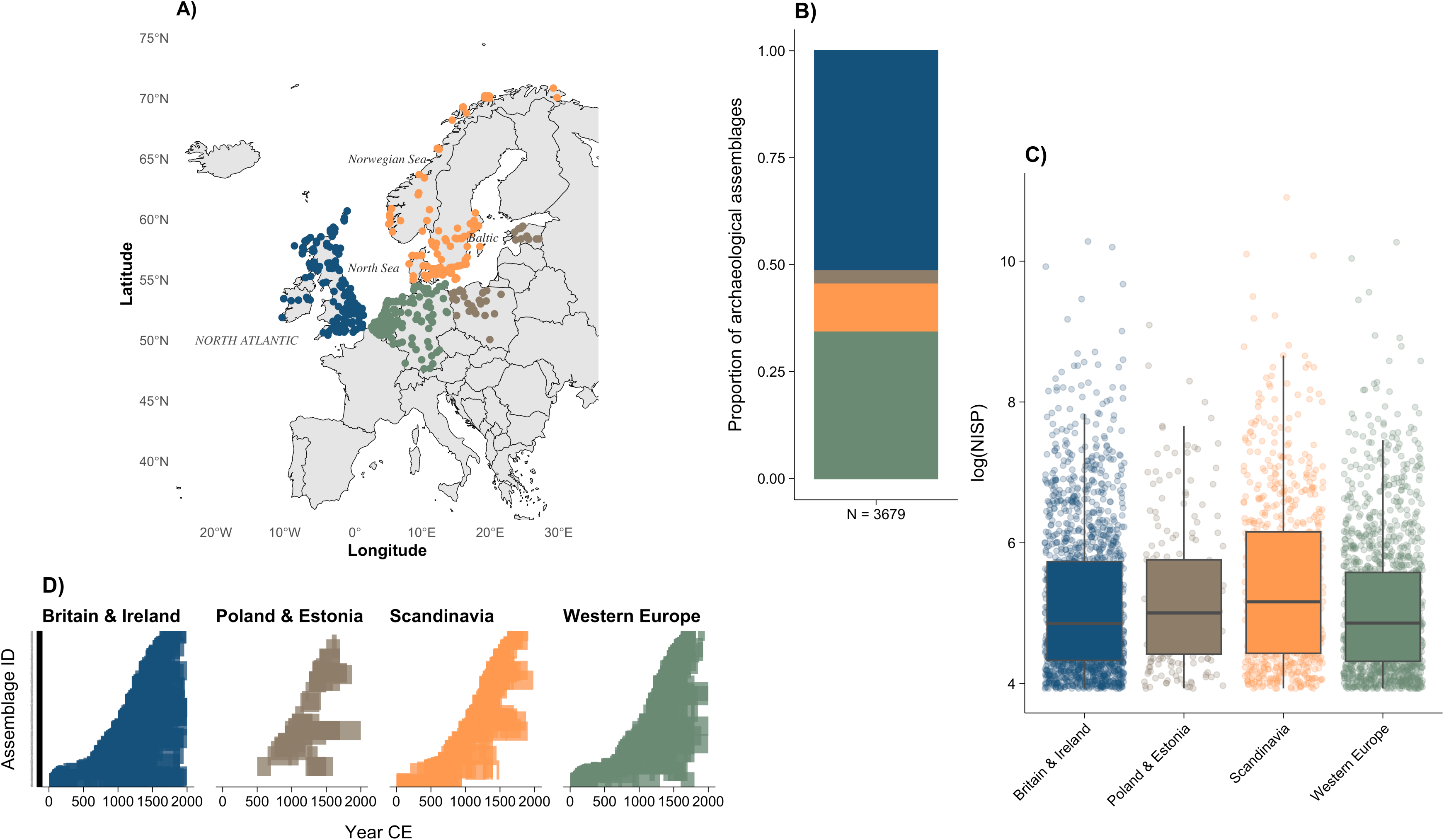
Summary of the NSS-DB. **a,** Geographic distribution of archaeological sites, coloured by region (Britain & Ireland (blue), Poland & Estonia (brown), Scandinavia (orange), and Western Europe (green)). **b,** Proportion of archaeological assemblages per region. **c,** Distribution of the number of identified specimens (NISP) recorded at assemblages within each region, on the log_10_ scale. **d,** Distribution of chronological ranges associated with assemblages in each region.

## Results

### Fish temperature proxy detects climate change during trade intensification period

Between 300–700 CE, Europe experienced pronounced cooling and heightened climate variability, driven in part by increased volcanic activity and reduced solar flux. In combination with socio-political factors, including the 5th-century collapse of the Western Roman Empire, these climatic shifts contributed to agricultural disruption, epidemics, famine, and mass migration^30,31^. We detected this environmental period, commonly termed the Dark Age Cold Period, using median temperature of catch (MTC) estimates (**Fig. 2**). While this period was less prominent in Western European and urban-associated assemblages (**Fig. S1**), a pattern emerged that was consistent with the possibility of lessened cooling in mainland Europe relative to the United Kingdom and Scandinavia. Notably, MTC suggests a pre-536 CE climate cooling, supporting existing evidence of a prolonged Dark Age Cold Period^31^ prior to the abrupt start of the so-called Late Antique Little Ice Age (c. 536 to 660 CE)^30^. MTC ranged variably between 7.3 and 10.8°C consistent with previous estimates of temperature variation in Europe (**Fig. 2**). MTC trends were significantly correlated with some previously published North Atlantic SST and precipitation rates^32,33^, although variability between proxies and differences in chronological resolution somewhat complicate such comparisons. During the second millennium CE, it is possible that faunal assemblages (and thus MTC) decreasingly reflect local temperature conditions due to a growing influence of trade^17,34,35^, homogenised culinary preferences^36–38^, and aquaculture^17^, leading to long-range transport of species and preferential selection of taxa (with particularly thermal tolerances). Nevertheless, cooling sea surface temperatures were observed during the MCA, consistent with proxy data from the Norwegian Sea, the subpolar North Atlantic, and North Africa^39–41^. Continuing cold MTC results were observed during the Little Ice Age (c. 1300 – 1850 CE)^27,42–44^, a period marked by harsher winters, crop failures, more frequent famine, and episodes of documented periods of political/social unrest^27,42,43^, indicating trade may not yet have significantly obscured temperature fluctuations.

**Figure 2.**
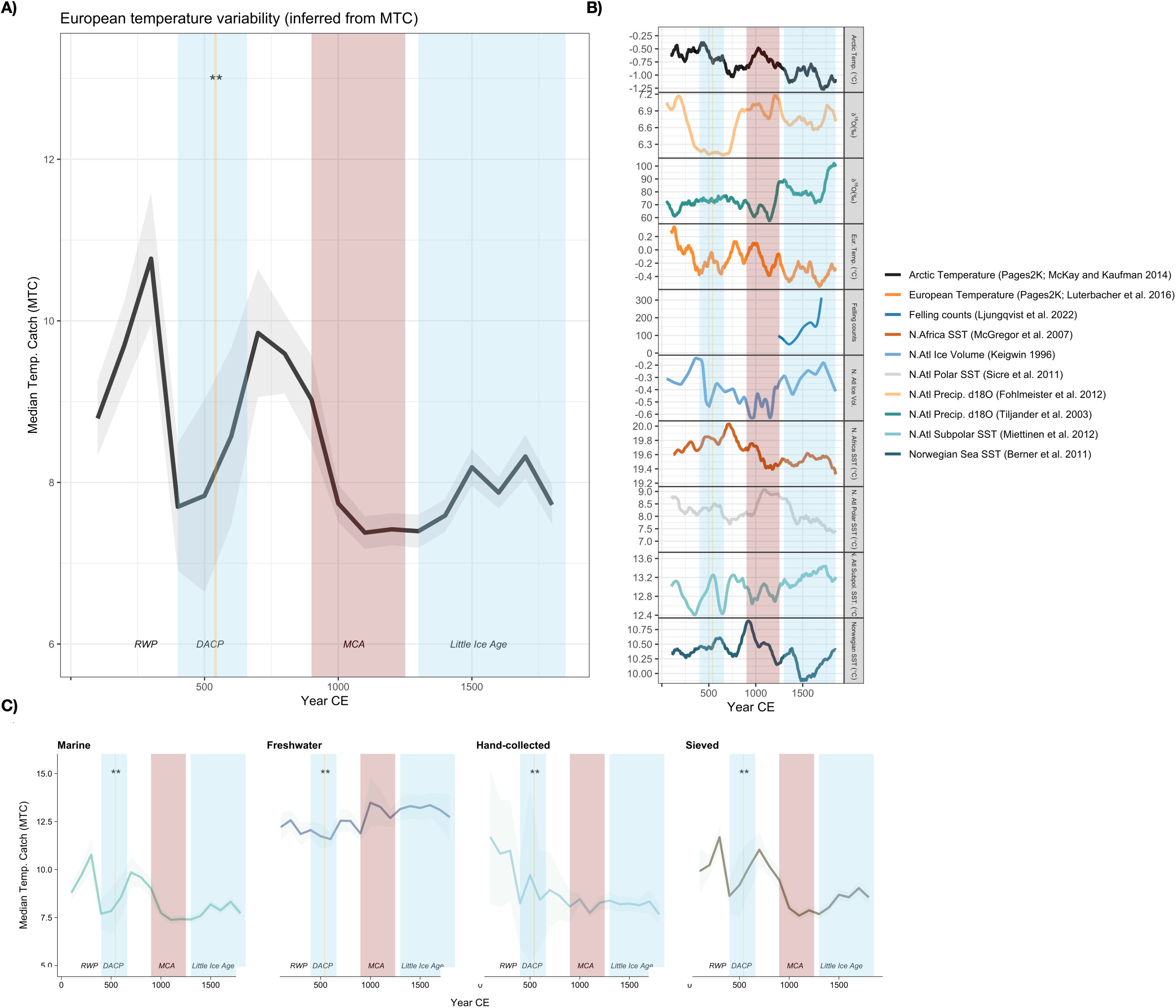
Median catch temperatures in relation to European environmental and demographic proxies. **a,** Time series of median temperature of the catch (MTC) calculated using thermal tolerances of fish species identified in NSS-DB (calculated using marine-species only). Mean (dark line) and standard deviation (shaded area) of MTC calculations across 1000 simulations. **b**, Comparative time series of selected environmental and demographic proxies from the literature, alongside NSS-DB MTC. **c,** Time series of sensitivity analysis for MTC shows freshwater do not reflect environmental proxies or NSS-DB MTC of marine species. From left to right: MTC calculated using marine species only, MTC calculated using freshwater species only (without known aquaculture sp.), MTC calculated using fauna from assemblages recovered by hand-collection only; MTC calculated using fauna from assemblages recovered by sieving only; this shows species from hand-collected assemblages do not reflect NSS-DB MTC temperature proxy from marine species.

Trade increased over time, with marked expansion of urban supply chains in the 11th to 13th centuries CE^15^, coinciding with the MCA, and of very large-scale trade networks during the 13th–16th centuries CE^17,45^, coinciding with the Little Ice Age. Trade expansion culminated in the establishment of global enterprises, including fisheries^8,46,47^. Between the 6th and 16th centuries CE, decreasing ecological dissimilarities of NSS-DB data suggest that catch compositions gradually homogenised at a cross-continental level (**Fig. 3**). This homogenisation was most marked between the 14th and 16th centuries, during the Little Ice Age (**Fig. 3**), suggesting that a combination of trade and increasingly shared dietary practices homogenised faunal deposits during centuries of heightened environmental challenges, demographic change and socio-ecological volatility.

**Figure 3.**
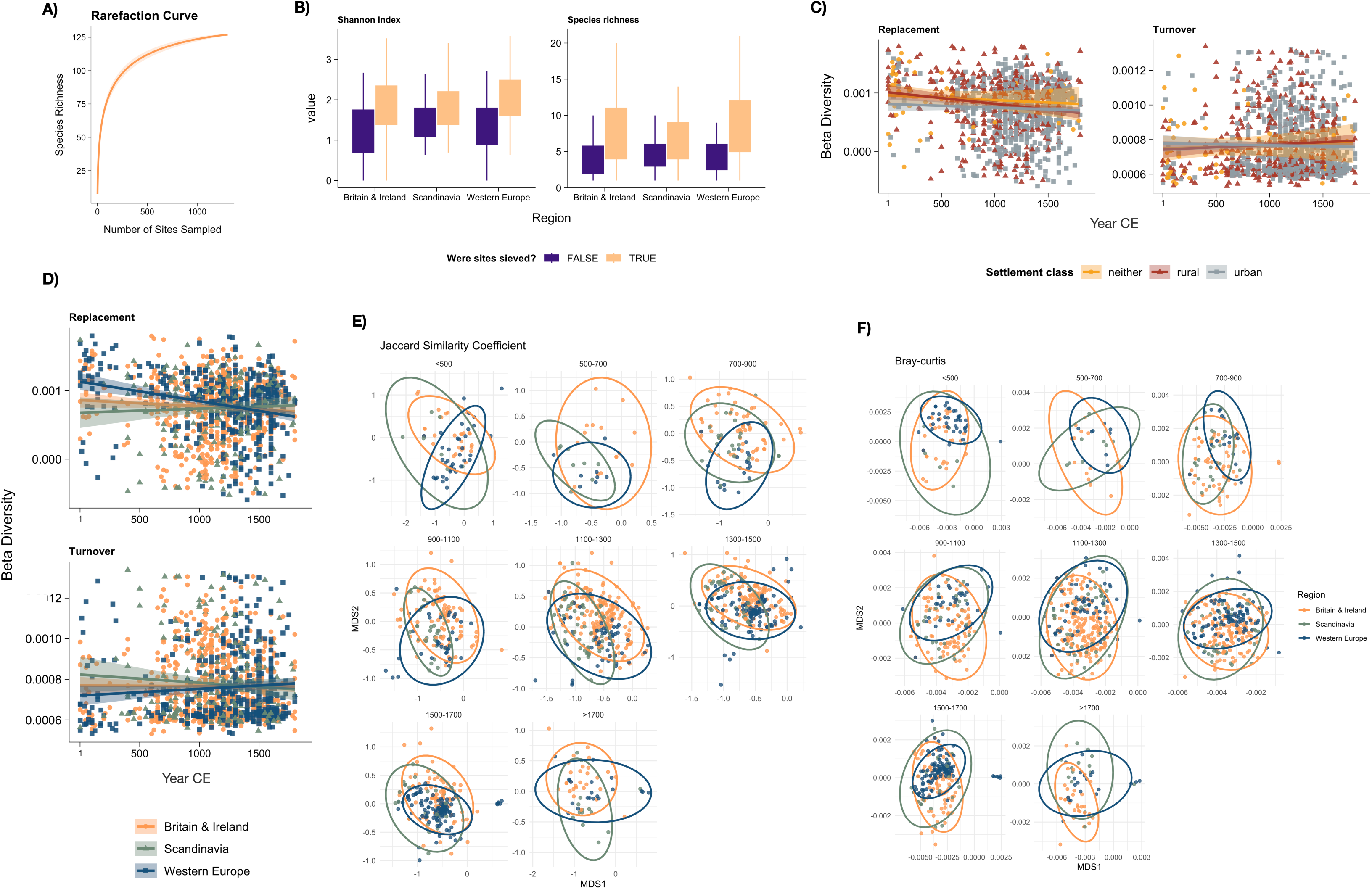
Temporal convergence and divergence in fish community composition revealed through diversity metrics and non-metric multidimensional scaling (NMDS). **a,** Expected increase in species richness with additional sampling of archaeological sites, shown as a rarefaction curve. Rarefaction was produced using 100 permutations using^110^. **b,** Boxplots of Shannon index and species richness in relation to region and whether sites were recovered using sieving. **c,** Temporal variation in beta diversity partitioned into replacement and turnover varied by settlement class (rural, urban, neither). **d,** Temporal variation in beta diversity partitioned into replacement and turnover varied by region. **e,** NMDS ordination of NSS-DB based on Jaccard Index disimilarities of NISP proportions identified to the species level. **f,** NMDS ordination of NSS-DB based on Bray-Curtis dissimilarities of NISP proportions identified to species level. Each point (**e,f**) represents an archaeological assemblage. Archaeological assemblages were aggregated by eight time bins and points coloured by region (Britain & Ireland, Scandinavia, Western Europe. Proximity of points indicates similarity in community composition and relative species abundances at those assemblages. Ellipses denote 95% confidence intervals around the centroid of each region’s assemblage distribution.

### Human diets increasingly relied on marine species

Faunal assemblages indicate an increase in marine fish consumption over time. In Western Europe and urban sites, the quantity of marine fish consumed homogenised trans-regionally by 1200–1400 CE, albeit slightly higher consumption in rural Britain & Ireland (**Fig. 4**). Overall, Scandinavia and Britain & Ireland had a significantly higher proportion of marine fishes relative to Western Europe (*p* < 0.05), potentially indicating higher marine fishing and/or consumption in these regions on average over the past two millennia (**Fig. 4**). By 1500 CE, all regions displayed comparable (0.5 – 0.75) proportions of marine fish, regardless of settlement class, indicative of a pan-regional homogenisation in the varieties of fish incorporated in human diets. Methodological differences in faunal recovery also influenced assemblage composition. Archaeological assemblages that had been sieved, at least as part of the recovery process, yielded a higher diversity of species (**Fig. 3(B)**) but a lower proportion of marine taxa (**Fig. 4(A)**), showing the importance of sieving in capturing smaller, predominantly freshwater species; despite this observation, reanalysis of MTC restricted to sieved assemblages supported previous temporal (**Fig. 2(C**)) and diet homogenisation patterns (results not shown).

**Figure 4.**
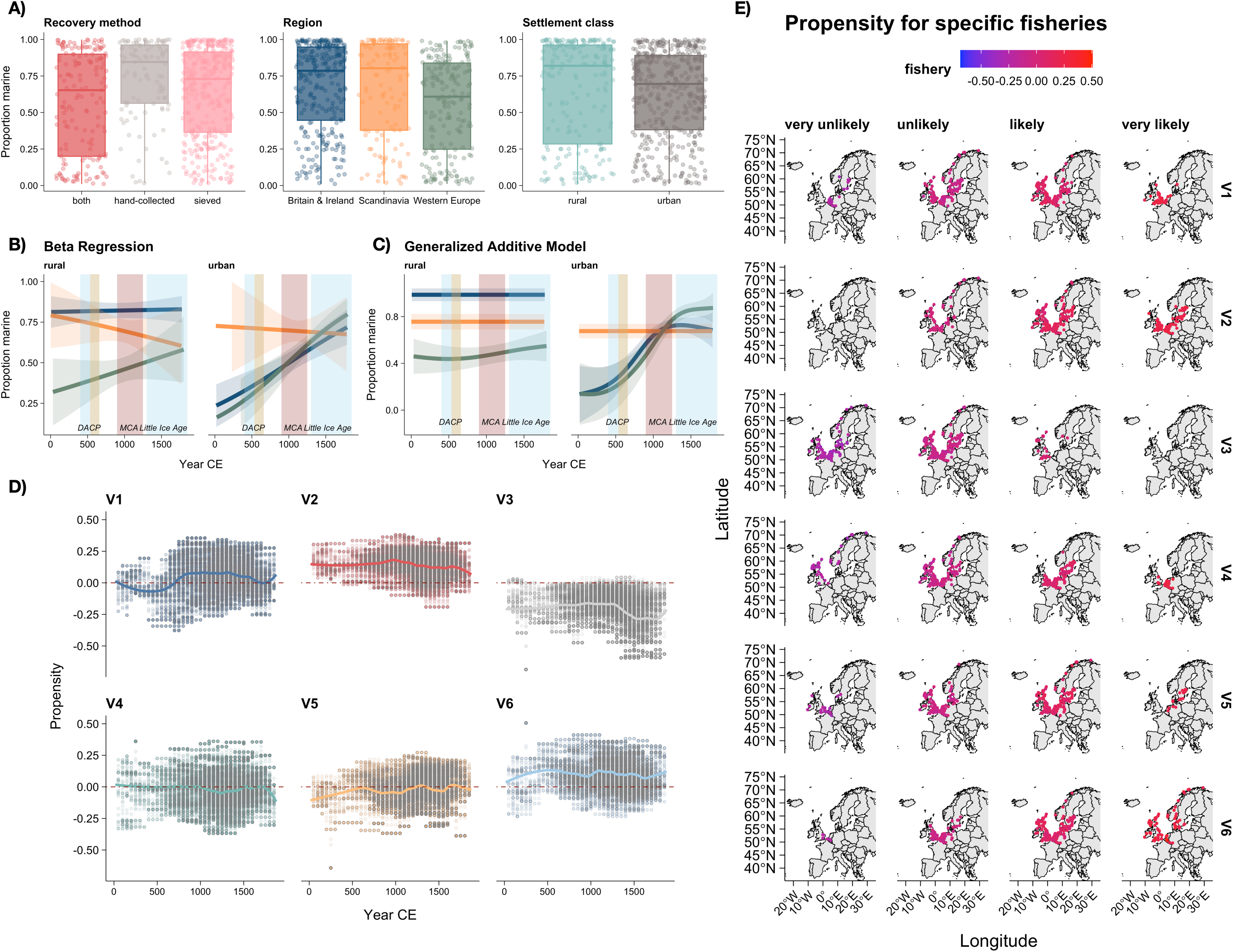
Temporal and spatial variation in European fishery propensity. **a,** Variation in the proportion of marine taxa by recovery method, region, and settlement class. **b,** Beta regression models showed that the proportion of marine species varied overtime and differed between region and settlement class. **c,** General additive models supported findings from beta regression. The best-fitting GAM model using the R package mgcv^118^. **d,** Temporal variation in the propensity of fishery topics (V1-V6) from recommender systems: V1-Marine-dominated fishery; V2-Mixed marine and freshwater fishery; V3-Not-commercially traded species; V4-Freshwater-dominated fishery; V5-Specialist marine and aquaculture fishery; V6-Deeper-water marine fishery. **e,** Spatial variation in the propensity of fishery topics shows broad geographical propensity. 0 = no propensity; < -0.2 is very unlikely; > 0.2 is very likely. Fisheries (V1-V6) as described in **d**.

Patterns of increasing marine harvesting were further corroborated by machine-learning models (recommender systems), which identified six recurrent fishery topics (**Table S1**). The first fishery (V1) was an obligate marine fishery that increased between 700–1000 CE and remained high thereafter until c.1700 CE. The second (V2), dominated by freshwater, inshore and estuarine species, exhibited relatively stable propensity from 1–1000 CE, followed by a gradual decrease thereafter. The third (V3) represented a diverse group of fish not likely to occur, with limited commercial importance, and thus were not considered further in analyses. The fourth (V4), characterised by freshwater fishes, showed relatively consistent propensity at approximately 50% of assemblages over time, with a decrease between 1000–1500 CE. The fifth (V5) was a mixed freshwater and marine fishery with slight peaks in the Dark Age Cold Period and the Little Ice Age. The freshwater taxa in fishery V5 are particularly associated with aquaculture, so the Little Ice Age peak may be related to an increased use of fish ponds in Europe at this time^17^. The final fishery (V6) was a deeper-water marine fishery that steadily increased between 1–550 CE, slightly declined following the Dark Age Cold Period and increased again more abruptly around 1000 CE at the beginning of the Medieval Climate Anomaly (**Fig. 4(E)**).

In addition, pairs of species with obligate marine life-histories (oceanodromous-oceanodromous) or obligate freshwater life-histories (potamodromous-potamodromous) increased between 500–1300CE relative to null expectations, with slight decreases thereafter, indicating a growing reliance on fish from specific habitats during these centuries (**Fig. S2**). In later time periods (1300 CE to post-1700 CE) mixed pairs became more apparent with a significant increase of oceanodromous-potamodromous pairs relative to null expectations (**Fig. S2**).

### Human diets increasingly relied on marine species: Transition to trans-regional diets

Local fish assemblages became less unique over time, demonstrating a gradual homogenisation until c.1500 CE (**Fig. 3**). This trend was not an artefact of species richness, which did not increase over time (**Fig. 3**). The pattern suggests a shift towards trans-regional fish dietary choices. Urban assemblages showed evidence that this homogenisation had already culminated by 1100–1300 CE, whereas rural sites did not follow suit until around 1300–1500 CE. Some convergence of fish dietary choices between Scandinavia and Britain & Ireland is also evident in the time bin 700–900 CE (**Fig. 3**), broadly concurrent with the early Viking Age when migration, settlement, and trade between these regions increased^14^.

As with NMDS results (**Fig. 3**), similarity tests showed species compositions at rural and urban sites became more similar over time, and similarities between regions dramatically increased (> 0.5) at urban sites from 900–1100 CE onwards and at rural sites from 1300–1500 CE. These shifts likely reflect the confluence of economic expansion, changing culinary preferences, and technological advancements in fishing practices (e.g. improved preservation and transport systems, aquaculture)^3,38,48^, that dominated urban areas and filtered into more rural regions thereafter. These trends highlight the emergence of trans-regional fisheries, alongside increased reliance on marine resources (**Fig.4(B**)) and the rise of freshwater aquaculture as essential parts of life especially during the Medieval Climate Anomaly and Little Ice Age.

Pairs of urban assemblages showed high rates of species similarity already from 900–1100 CE, with c. 80% of pairs showing greater similarity than null expectations (results not shown). In contrast, rural-rural pairs showed a gradual increase in similarity from c. 60% during 900–1100 CE, increasing steadily thereafter (c. 70% in 1100-1300 CE, and exceeding 80% by 1500-1700 CE), a trend mirrored in rural-urban pairs. These patterns suggest diet homogenisation began earlier in urban centres, with rural and urban assemblages becoming increasingly alike over time. We infer that the standardisation of marine fish consumption practices spread to rural areas during the late Middle Ages and into the early modern period, coinciding with the expansion of trade networks, market integration, increased human mobility, and the convergence of cultural practices^49^; this period included the emergence of super-regional trade organisations, such as the Hanseatic League^49^. Both homogenisation analyses (**Fig. 3**) and similarity tests suggest culinary practices began to re-diverge by 1700 CE towards the end of the Little Ice Age.

The Little Ice Age was characterised in much of Europe by glacial expansion, long and severe winters, and more frequent harvest failure^26,27,50,51^. Climatic stressors occurred in tandem with dramatic demographic changes, including population collapse due to 14th-century famine and plague in parts of the continent^51^, followed by subsequent recovery (**Fig. 2(B)**). Reduced agricultural production associated with climatic cooling, coupled with labour shortages and rising real incomes after the Black Death^51^ may have contributed to shifting dietary patterns. One possible outcome was an increased demand for alternative protein sources, reflected in the continued expansion of marine fish consumption that we observed in Western Europe at the transition from the Medieval Climate Anomaly to the Little Ice Age (**Fig. 4**). However, this was likely one of several factors shaping demand: dairy, grain, and meat would also have played a role^52,53^. Moreover, the developments in these marine fisheries may have influenced availability and affordability. Increased marine fishing may have resulted in a surplus of stockfish, lowering its price relative to other protein sources (e.g. freshwater fish, meat). Historically observed expansions in marine fishing, due in part to the opening of the North American fishing grounds during the late 15th to early 16th centuries, resulted in a stockfish surplus in Western European markets^8,37^. Declines in price for cod/stockfish, both in real value and in relation to inflation, were reported in England, the Netherlands, France, Germany and Norway from the late 15th to late 17th centuries^37,48,54^.

### The Northern Seas Synthesis Database (NSS-DB)

Fisheries have played a pivotal role in shaping social, economic, and political change in Europe^3,17,55^. The Northern Seas Database (NSS-DB) captures two millennia of ichthyoarchaeological data, documenting the relationship between fishing and fish consumption. It contains 1,910,794 identified fish specimens from 282 taxa, spanning 30 orders, 66 families, and 158 species, many of which are vulnerable to extinction today^56^. By systematising legacy data, the NSS-DB enables quantitative historical ecological research, overcoming limitations that often confined past analyses to qualitative assessments, or localised, often taxa-specific case studies^2,57,58^. Here, we demonstrate its potential for investigating climate change, trade, aquaculture, and shifting culinary practices, all key natural and cultural dynamics known to shape past societies for millennia^4,6,9,17,59^. Openly available on Dryad, the database now offers a powerful tool for researchers to explore long-term changes in fishing practices, shedding light on how extraction, management, and habitat shifts influenced species demographics, local extirpations, and modern extinction risks^56,60,61^ across northern and Western Europe. Future integration with biomolecular research (e.g.^62^), stable isotope analysis^15^, and historical and palaeoecological evidence^63^, will enhance our understanding of past fishing pressures, management strategies, and climatic changes, at local, regional, and continental scales. Beyond that, the NSS-DB complements analogous research across Europe and the wider Northern Hemisphere, unlocking new opportunities to assess the long-term interplay between fisheries, ecosystems, and connectivity between human societies further afield^64^.

## Discussion

Our trans-regional European synthesis of archaeological fish remains offers quantitative evidence linking environmental volatility with shifts in fishing practices, trade, and dietary homogenisation across Europe over the past two millennia. Using the NSS-DB, we detect key climate trends, such as the Dark Age Cold Period^31^ and the Little Ice Age^27^. We also detect the counter-intuitive cooling of sea surface temperatures during the Medieval Climate Anomaly^39–41^, before the onset of the Little Ice Age as inferred from terrestrial air temperature proxies^30,65^, a pattern that is consistent with growing knowledge regarding the complex relationship between sea surface temperatures and atmospheric circulation^66^. We infer important socio-economic changes associated with the Medieval Climate Anomaly (commonly dated 950 – 1250 CE, but recognised as having regionally variable onset and duration, with some warming beginning as early as 800 CE) and the Little Ice Age^27,42,43,65,67^, such as expanding trade networks and dietary homogenisation. Trade networks can emerge in response to environmental stress, enabling the acquisition of distant resources and technological innovations^9,10,35,68^. Here we see the requirements of rising urbanism and population growth mitigated by increased marine fishing during the Medieval Climate Anomaly, and the stresses of climatic volatility and plague mitigated by continued and expanded marine fishing, and perhaps also by aquaculture during the Little Ice Age (**Fig. 4**). These developments coincided with homogenising culinary practices (**Fig. 3**), starting in towns and spreading to the countryside, suggesting growing reliance on long-distance trade. Together, these trends align with wider archaeological and historical evidence for expanding European trade networks, alongside demographic growth, rising technological capabilities and preservation methods (e.g. the Dutch herring buss^69^; increased solar salt production^37^), and shifting culinary practices^37^. Collectively, our findings support research that trade networks and culinary innovations often expand in response to environmental change and instability^9,10,35,68^.

Urban and rural centres experienced transformations of marine fishing in distinct ways. Rural sites (especially when coastal) had access to marine fish throughout the last two millennia, while urban centres showed increasing consumption of obligate marine taxa through time (**Fig. 4**), with a further emphasis on deeper water and traded sea fish emerging during the MCA (**Fig. 4, V1, V6**). Conversely, trans-regional dietary homogenisation occurred later at rural sites. In urban contexts, increased reliance on marine fisheries likely reflects the combined effects of rising population densities (both preceding and following the demographic crash associated with the Black Death), reduced agricultural productivity in parts of Europe during the Little Ice Age, increased purchasing power in the late Middle Ages and also mandated abstinence of meat consumption on Fridays, during Lent and other pious days due to medieval Roman Catholic practices (prior to the 16th-century Reformation for affected regions)^70,71^. By the 12th century CE onward, similar proportions of obligate marine fish were observed at rural and urban sites (**Fig. 4)**. Marine fish continued to be important at both urban and rural sites in later centuries. These patterns differ somewhat from previously observed trends in the eastern Baltic, where marine fishing increased later, from c.1200 CE onwards, and to a lesser degree relative to Western European sites^72,73^.

The 1500s marked the beginning of what has been termed the ‘North Atlantic Fish Revolution’^8,37^ , during which regions such as Norway, Iceland and Newfoundland emerged as pivotal suppliers of dried and salted cod and other preserved fish, and the Netherlands a key supplier of herring, for European markets^37^. This expansion may be reflected in the latest time bins of our composition analyses, where some inter-regional divergence is observed from the 1500s onwards (**Fig. 3**), following the preceding convergence of the Middle Ages (**Fig. 3**). Such transitions are often challenging to identify in the archaeological record, as postmedieval zooarchaeology in Europe has received comparatively little research interest^74^, and archaeological finds from recent centuries are not always protected under national laws (e.g. non-Sámi terrestrial sites post-dating the 1537 Reformation in Norway). Moreover, the method used for recovering archaeological material may not always capture the entire community composition of fauna due to missing species. Nevertheless, shifts towards trans-regional markets^75^, national religious reformations^76^, and the expansion of global maritime empires^77^ were so profound that their signal has been captured in aquatic faunal assemblages across northern and Western Europe, and now illuminates a fundamental restructuring of human-aquatic interactions.

The fishery homogenisation observed in our data mirrors well-documented processes in other aspects of food production and land-cover change^78^. Similar homogenisation trends have been observed in crop composition^79,80^ and commonly-kept livestock^9,81^. Just as ancient grains and local landraces have been supplanted by high-yield, globally distributed varieties, regional fish assemblages were supplemented by a core set of widely traded marine taxa. Homogenisation of culinary practices^36,82^ can also be attributed to enhanced market integration and trade expansion (e.g. the German Hanseatic League and its antecedents, from the late 12th century onwards ^49^). Importantly, this cultural shift focused demand to such a degree that even once superabundant fish stocks such as Baltic autumn-spawning herring^62^ and Newfoundland’s Northern cod^83,84^ collapsed under sustained harvesting – supporting that diet diversification may be vital to sustainable human resource management^85^.

Our results draw on a combination of classical statistical techniques as well as the newest methodological developments in their application, including recommender systems of machine learning. The use of multiple complementary methodologies (termed methodological omnivory for historical sciences^86^) strengthens the plausibility of the results. Our findings support a wealth of research highlighting the long-term dependence of humans on aquatic resources^87,88^. The ever-expanding, often overconsumption of oceanic resources has resulted in the depletion of marine fish stocks worldwide^89^. Conservation efforts are ongoing to restore fish populations^90,91^, however, many species remain depleted and are critically endangered or threatened today^92^. The NSS-DB houses two millennia of information on the relationship between fishing, fish and fish consumption, thereby providing a tool to test how past human pressures may have shaped fish demographics and extinction statuses in the modern day. Great potential exists to link this dataset with the results of analogous research covering other areas and time periods in Europe and across the Northern Hemisphere (e.g.^64,93^). Moreover, through comparison with other evidence (e.g. biomolecular, historical written sources), the dataset can help to explain how specific fisheries have been of paramount importance to past social, economic and political change in Europe^14,17,57^. The world’s oceans are undergoing rapid change^94,95^, and global decisions on extraction and conservation are being made annually^96^, thus addressing shifting baselines with a wealth of historical data, such as the NSS DB, is more paramount than ever.

## Methods

### Database curation

Fish bone counts with known origin and time period dating between 1 CE and 2000 CE were contributed by experts (see Author Contributions). Data were provided via a Microsoft Excel template with an accompanying data dictionary, or through published or unpublished reports. Only publicly available or authorised data were included, with intellectual property status provided. Counts of fish bones (number of identified specimens, NISP) were recorded across 11 countries at the level of archaeological assemblage, defined primarily by a locality with a given time range (chronology) and known recovery method (e.g., sieved, hand-collected); regional stratigraphic resolution of assemblages varied between datasets. Additional metadata, such as precise chronology and standardised georeferences were supplemented by sourcing excavation reports, or using Google Earth or Google Maps. To evaluate the potential error rate of data-entry, 5% of the assemblages in the England dataset were recompiled (error rate ∼10%).

Fish taxa were retained as reported by data contributors. Typographical errors, variation in nomenclature, and use of common names were reconciled post-data collection using the R package ‘taxize’^97^. Taxonomic backbones were obtained also using taxize and rGBIF^97,98^ and taxon assignments manually checked. Corresponding NISP data for any remaining taxonomic uncertainties were excluded. For taxa reconciled at the species level, ecological trait data (i.e. life history, temperature) were obtained using^99^.

The database (NSS-DB v1) contains 1,910,794 NISP from 3679 archaeological assemblages comprising 30 orders, 66 families, and 158 species. Data originate from four broad regions: Britain & Ireland, Scandinavia, Western Europe, and Poland & Estonia (**Fig. 1**); these designations are descriptive, reflecting incomplete regional coverage and the potential for future database expansion (e.g. the inclusion of more Western European or Mediterranean assemblages). Data were cleaned and aggregated to the species-level, assemblages with less than five zooarchaeological specimens and assemblages in the top 2.5% of species-level specimen counts were excluded to minimise potential impact of outliers due to sampling biases. Assemblages from Poland & Estonia were removed as the majority of these sites were not recovered using sieving and this is likely to bias likelihoods of species recovery, resulting in 621,423 NISP across 1305 assemblages for analysis.

### Median temperature of the catch (aMTC)

Thermal tolerances of fish species (identified in the NSS-DB) were used to track temperature fluctuations across Europe over the last two millennia. Median temperature of the catch (MTC) analyses were estimated across NSS-DB using data from three of the four regions (Britain & Ireland, Scandinavia, Western Europe, see **Fig. 1**). MTC calculations were also calculated independently for each of the three regions. Median, minimum and maximum temperature preferences for each species were obtained from FISHBASE^100^ following^25^. When observed temperature ranges were unavailable, modelled temperature preferences were extracted (available from FISHBASE^100^ using aquamap estimates^101,102^). Fish with temperature range distributions in the top 5% (c.>15°C) were excluded as these species were unlikely to provide information on local environmental conditions.

To address chronological uncertainty of archaeological assemblages, each assemblage was randomly assigned to a 100-year time bin within their specific chronological range; temperature calculations were run 1000 times, with replacement, providing uncertainty in temperature fluctuations associated with chronological uncertainty.

Median temperature of the catch was estimated for each 100-yr time bin using the equation from^25^ (using NISP rather than MNI), originally adapted from^103^ as follows:

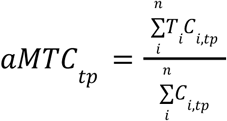

*Ci,tp* is the total number of specimens identified for a given species (*i*) in a particular time period (*tp*). *Ti* is the median temperature preference of a species (*i*), and *n* is the total number of species. This equation weights the proportion of a given species within a specific time period by multiplying the median temperature preference of each species by the number of identified specimens and dividing by the total count for that time period. For sensitivity of temperature trends to median values, minimum, and maximum temperatures were additionally used in replacement of *Ti*.

Changes in temperature over time were assessed using the autocorrelation function *acf*^104^. Correlations between aMTC results and existing temperature proxies (**Fig. 2**) were assessed using cross-correlation (function *ccf*) using the R package ‘stats’^104^.

### Trans-regional European trends in fish consumption

To assess how the diversity of fishes in human diets changed over time, the number and evenness of fish taxa consumed were assessed by measuring alpha and beta diversity of fish taxa identified in archaeological assemblages^105–107^. Chronological uncertainty and time-bin sizes were mitigated via randomised allocation as above (1000 simulations, 100-yr bins). Similarity in taxonomic composition of the catch (species-level counts from NSS-DB) across eight time periods (< 500 CE, 500–700 CE, 700–900 CE, 900–1100 CE, 1100–1300 CE, 1300–1500 CE, 1500–1700 CE, > 1700 CE) was assessed using two approaches: pairwise ordination (NMDS) following^108^ and recommender systems (adapted from^109^). For ordination, Bray-Curtis (weighted by counts) and Jaccard (presence/absence only) indices were estimated using the R package ‘vegan’^110^ and resulting indices inverted so that values closest to 1 and 0 represent most similar and least similar, respectively.

To explore whether fishing and fish consumption (collectively hereafter as ‘fishing practices’) became more homogenised over time, pairwise similarity tests were performed using an assemblage-based taxonomic-occurrence matrix (structured with archaeological assemblages as rows and taxa as columns and with 0’s marking taxon absence and 1’s marking taxon presence at sites). Cosine similarities were calculated (following^109^) for each assemblage whereby pairwise similarities of two assemblages were a function of the co-occurrences of the taxa at each of those assemblages. Average similarities in pairs of rural and urban sites within or between regions were calculated over time, under the hypothesis that rural or urban sites within a given region were more likely to be similar to one another in earlier time periods compared with later ones, as cultural transmission increased and fishing practices became homogenised between regions. Recommender systems are models trained using machine-learning algorithms commonly used in e-commerce and online advertising to enhance user experience^111,112^. The capabilities of these models for analysis of ecological datasets are only recently being realised, adapted as a form of joint species distribution modelling and for estimation of species abundances^109^. Recommender systems are designed to work well with sparse datasets, where a high proportion of input data are zeroes and there is high uncertainty in those recorded absences^113^. The models capture patterns in co-occurrences of taxa and use them to predict the propensity of taxa to occur where they have not been observed, making them an ideal candidate for analysing ecological patterns in the past from archaeological or paleontological assemblages where taphonomic processes degrade the record relative to true living assemblages^109^. Here, we use recommender systems to identify key fisheries (fishing and fish consumption combined) that have been consistently observed across Europe over the past two millennia and identify any preferential shifts towards freshwater and/or marine fishing.

Implementing recommender systems using a matrix factorization method predicts propensity scores for each assemblage and taxon to occur in a given fishery (the term ‘fishery’ refers to each column in the inner matrices produced from iterative singular value decomposition of the input matrix^114^). Inner matrices were calculated using matrix factorization, based on the number of archaeological assemblages and taxa, while inner dimensions were evaluated with neighbourhood sizes (K) of 3,4,5,6,7,9, and 12. Model performance was assessed using the area under receiver operating characteristics curve (AUC)^115^. After evaluating recommender system model performance for a range of parameters, the best fit model under AUC contained six fisheries (V1-V6) that exemplify patterns in fish use by human societies across Europe. Fish species were characterised as known major traded taxa or not. We assess these fisheries’ histories using nonlinear time series and Generalized Additive Models (GAMs). Propensity of fisheries were modelled as a function of region (Britain & Ireland, Scandinavia, Western Europe), time (100-yr intervals), recovery method (hand-collected, sieved, both), settlement class (rural, urban, neither), urban centre (boolean), total NISP, and chronological range using the R package ‘mgcv’. Model performance was assessed using Akaike’s Information Criterion (AIC)^116^.

Finally, to identify whether a preference for marine fishing increased over time two methods were used. First, beta regression and GAMs were used to identify temporal variation in the proportion of obligate marine species within archaeological assemblages using the R packages ‘glmmTMB’ and ‘mgcv’, respectively^117,118^. Year, chronological range, recovery method, region, and settlement class were included as fixed explanatory variables and two-way interactions. Model selection was based on weighted-AIC. Comparisons between fit of beta regression and GAM models were assessed using the deviance explained from Adjusted R-squared (beta regression) and deviance explained in the best fitting GAM model.

Second, z-scores of cosine similarities from input matrices (no. assemblages X no. species) were compared relative to a null model. Null models shuffle taxon occurrences between assemblages 500 times and recalculate pairwise similarity. *z* > 1.96 indicates that it is unlikely that the observed similarity occurred at random (less than 5% probability). Average z-scores were calculated for one of eight time-bins (< 500 CE, 500–700 CE, 700–900 CE, 900–1100 CE, 1100–1300 CE, 1300–1500 CE, 1500–1700 CE, > 1700 CE) using: (i) pairs of obligate marine species; (ii) pairs of obligate freshwater species; (iii) mixed marine-freshwater pairs; or (iv) other (catadromous/anadromous). We inferred a higher ratio of significant marine pairs between time bins to indicate an increased reliance on marine species within that given time period.

## Data availability

The Northern Seas Database V1.0 (NSS-DB) can be downloaded online from Dryad (https://doi.org/10.5061/dryad.XXXX).

## Code availability

All code used to conduct analyses for this manuscript are available at the following links: https://github.com/DannyLBuss/Northern-Seas-Synthesis and https://github.com/abbaparker/NSS-Recommender-Systems. Figures 1, 2 ,3 and 4 were produced in RStudio version 2022.7.2.576 (R version 4.2.^119^).

## Acknowledgements

The zooarchaeologists responsible for collecting and identifying the material analysed herein are gratefully acknowledged and individually named in the NSS-DB files. The authors thank Alison Locker and Dirk Heinrich for their especially extensive data contributions and informative discussions throughout the project. This research has benefited from wider discussion within the 4-OCEANS project team, especially including Cristina Brito, John Nicholls, Eva Jobbová, Francis Ludlow, and Bastiaan Star. The authors thank Michael Puma, Al Matthews, Katrien Dierickx, Jo Sindre P. Eidshaug, Wendy Hlengiwe Khumalo, Erin Kunisch, Bente Phillipsen, Mike Martin, Matt Page, Paxti Perez Ramallo, Martin Seiler, Eirik Sollid, and Youri van den Hurk for their collegial discussions and perspectives during the development of this work.

## Author contributions

**Conceptualisation**: D.L.B. and J.H.B. **Methodology**: D.L.B., A.K.P., I.Ž., and J.H.B. **Software**: A.K.P. and I.Ž. **Validation**: D.L.B., A.K.P., M.N.E., M.F.-A., I.Ž., and J.H.B. **Formal analysis**: D.L.B., A.K.P., and R.C. **Investigation**: D.L.B., M.F.-A., T.C.A.R., R.B., A.B., M.K.D., M.N.E., I.B.E., A.E., S.H.D., J.F.H., R.C.H., A.K.H., I.v.d.J., B.K., H.C.K., F.C.L., L.L., O.M., D.M., E.M., H.J.M.M., W.F.M., R.N., L.Q., H.R., K.R., A.S., W.V.N., W.W., and J.H.B. **Resources**: all authors. **Data curation**: D.L.B., M.F.-A., R.C., A.B., M.K.D., A.E., J.F.H., R.C.H., A.K.H., I.v.d.J., B.K., H.C.K., F.C.L., L.L., O.M., D.M., E.M., H.J.M.M., K.R., A.S., W.V.N., W.W., and J.H.B. **Writing – original draft**: D.L.B. **Writing – review & editing**: all authors. **Visualisation**: D.L.B. **Supervision**: J.H.B. and D.L.B. **Project administration**: J.H.B. and D.L.B. **Funding acquisition**: J.H.B. and P.H.

## Supplementary Figures

Fig. S1: Figure showing A) Time series of regional median temperature of the catch (MTC) across NSS-DB fish assemblages gathered from either rural or urban sites within Britain & Ireland (Blue lines), Scandinavia (Yellow lines) or Western Europe (Green lines). MTC values were calculated using NISP data for fish species grouped into 100-year time bins. The dark line represents the mean MTC across 1000 simulations, while the light blue shading indicates the standard deviation of MTC from the 1000 simulations. B) Time series of environmental proxies re-shown for comparison.

Fig. S2: For categories describing species pairs based on their life history categories, the proportion of pairs whose occurrence similarities in each time bin are significantly higher than in the distribution of cosine similarities of 500 null matrices. “Fresh” indicates pairs where both species are potamodromous. “Marine” indicates pairs where both species are oceanodromous. “FreshMarine” indicates pairs where one species is potamodromous and one species is oceandromous, so the two species are unlikely to be found in the same water body. “Other” indicates pairs where one of the species has any other life history category, including catadromous, anadromous, mixed, or unknown. Time bin numbers: 1= before 500, 2= 500-700, 3=700-900, 4= 900-1100, 5= 1100-1300, 6= 1300-1500, 7= 1500-1700, 8=after 1700.

## Supplementary Tables

**Table S1**. Results of fishery topics categorisation using species propensities and ecological traits

