## Supplementary figures and images for "2000-year fish bone record reveals transition to commercial fisheries during climatic change"

### Fig. S1

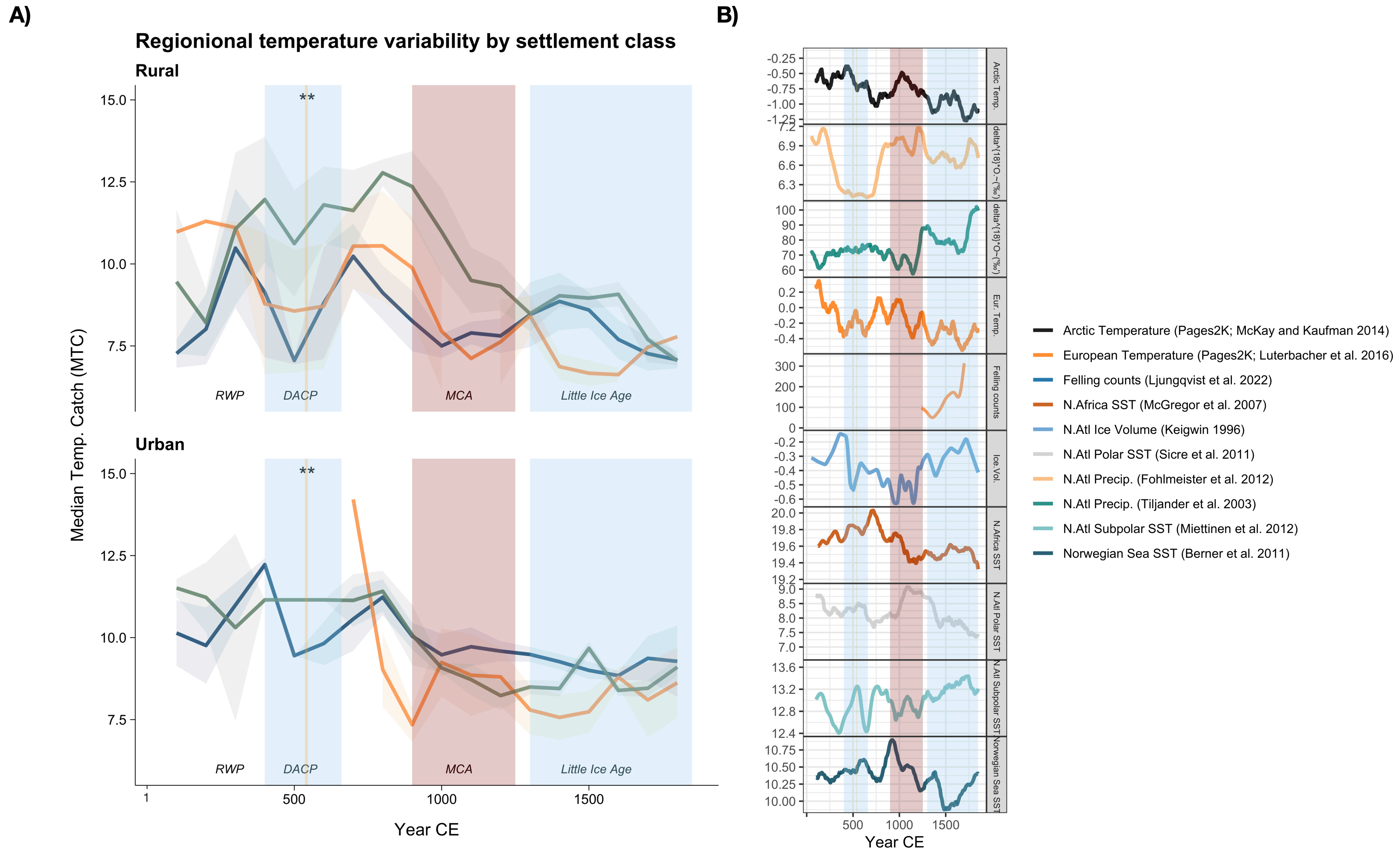

### Fig. S2

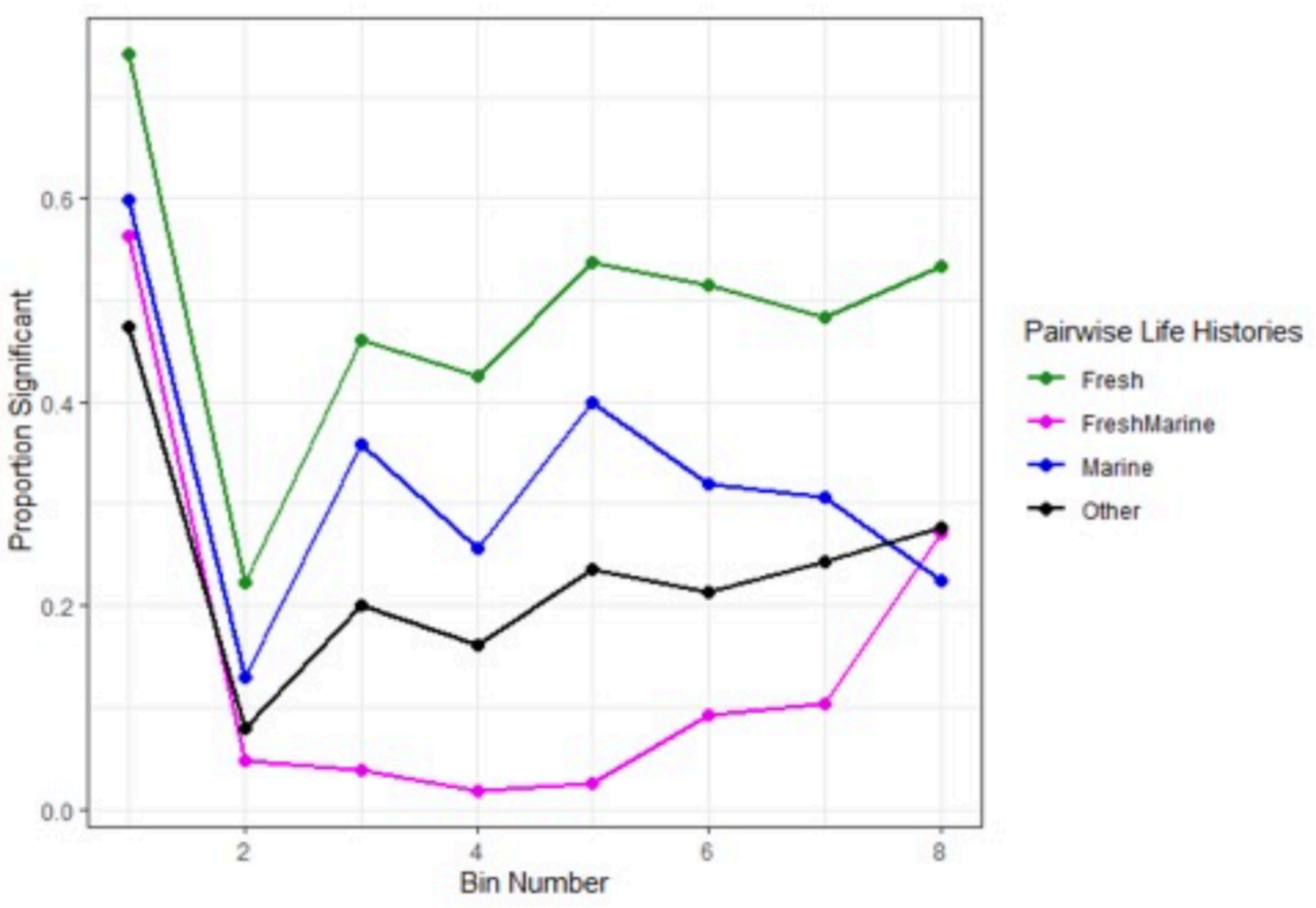
